# biMM: Efficient estimation of genetic variances and covariances for cohorts with high-dimensional phenotype measurements

**DOI:** 10.1101/087932

**Authors:** Matti Pirinen, Christian Benner, Pekka Marttinen, Marjo-Riitta Järvelin, Manuel A. Rivas, Samuli Ripatti

**Affiliations:** Institute for Molecular Medicine Finland (FIMM), University of Helsinki, Helsinki, Finland.; Helsinki Institute for Information Technology HIIT and Department of Mathematics and Statistics, University of Helsinki, Helsinki, Finland.; Department of Public Health, University of Helsinki, Helsinki, Finland.; Helsinki Institute for Information Technology HIIT and Department of Computer Science, Aalto University, Espoo, Finland.; Biocenter Oulu, University of Oulu, Oulu, Finland.; Department of Epidemiology and Biostatistics, MRC-PHE Centre for Environment and Health, School of Public Health, Imperial College London, London, UK.; Center for Life Course and Systems Epidemiology, Faculty of Medicine, University of Oulu, Oulu, Finland.; Unit of Primary Care, Oulu University Hospital, Oulu, Finland.; Department of Biomedical Data Science, Stanford University, Stanford, CA, USA.

## Abstract

Genetic research utilizes a decomposition of trait variances and covariances into genetic and environmental parts. Our software package biMM is a computationally efficient implementation of a bivariate linear mixed model for settings where hundreds of traits have been measured on partially overlapping sets of individuals.

**Availability:** Implementation in R freely available at www.iki.fi/mpirinen.

## 1 Introduction

Decomposing phenotypic variance and covariance into genetic and environmental parts is important for designing genetic studies and understanding relationships between traits and diseases. The two main approaches are linear mixed model (LMM) implementations, such as GCTA (Yang *et al.*, 2011), GEMMA (Zhou and Stephens, 2014) or BOLT-REML (Loh *et al.*, 2015), and LD-score regression, implemented in LDSC (Bulik-Sullivan *et al.*, 2015). LMM requires access to the individual-level genotype-phenotype data whereas LDSC only needs output from a genome-wide association study (GWAS) and variant correlations from a reference database, but consequently may be less precise than LMM (Bulik-Sullivan *et al.*, 2015).

We consider settings where individual-level data are available, and hence use LMM. The bivariate LMM for *n* individuals is 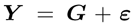, where 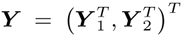 is 2*n*-vector of mean-centered phenotype values from which the covariates, such as age, sex and principal components of population structure have been regressed out, 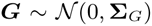 is 2n-vector of genetic random effects and 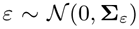 is 2*n*-vector of environmental random effects. The (2*n*) × (2*n*) covariance structures are parameterized by six scalars: genetic variances *V*_*G1*_ and *V*_*G2*_, genetic covariance *V*_*G12*_, environmental variances *V*_*ε*1_ and *V*_*ε*2_ and environmental covariance *V*_*ε*12_ as

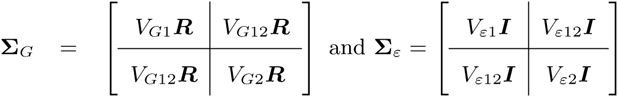

expressed as *n* × *n* block matrices. ***I*** is the identity matrix and the element *i, j* of the genetic relationship matrix (GRM) ***R*** is

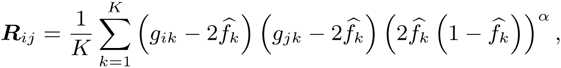

where *g*_*ik*_ is the genotype of individual *i* at variant *k*, coded as 0, 1 or 2 copies of the minor allele and 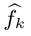 is the minor allele frequency (MAF). We use the standard scaling of allelic effects determined by *α* = −1.

From this model, an estimate of *V*_*Gt*_ approximates additive genetic variance of each trait (*t* = 1, 2) explained by the variants included in the calculation of ***R*** and is often used as a lower bound for the (narrow-sense) heritability (detailed assumptions in Yang *et al.* (2015)). An estimate of the genetic correlation 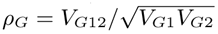 measures (average) correlation of the allelic effects of the variants on the two traits. Similarly, we can estimate 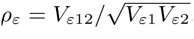, the correlation in the environmental components between the traits.

A challenge with bivariate LMMs, that operate on an explicit ***R*** matrix (e.g. GCTA and GEMMA), is that they require matrix operations cubic in the cohort size for each pair of traits analyzed, which becomes computationally prohibitive for handling hundreds of traits measured on 10,000s of individuals. Our software package biMM speeds up the bivariate LMM analysis (1) by a fast likelihood computation, (2) by reusing matrix decompositions across pairs of traits, and (3) by arranging data to optimize sample overlap between consecutive pairs of traits.

## 2 Methods

### 2.1 Reusing eigendecomposition

Once an eigendecomposition of ***R*** is available, our biMM algorithm drops the time complexity from cubic to quadratic for a trait pair and from cubic to linear for a single evaluation of the likelihood function (Supplementary Information). A crucial observation is that a complete sample overlap between two trait pairs means that the same eigendecomposition can be used for both pairs.

### 2.2 Ordering pairs, imputing and dropping values

We order the trait pairs in such a way that the consecutive pairs have a large sample overlap. biMM further allows imputing at most *t*_*i*_ missing values and/or dropping at most *t*_*d*_ non-missing values for a trait pair to make it match the available eigendecomposition (Supplementary Information). Only when this is not possible for any remaining pair does biMM a new eigendecomposition. Algorithmically, given user-specified *t*_*i*_ and *t*_*d*_, biMM finds an ordering that results in a small number of total eigendecompositions. This is an instance of the shortest Hamiltonian path problem that we tackle by a greedy heuristic (Supplementary Information).

### 2.3 Example analysis

We consider data from the Northern Finland Birth Cohort 1966 (NFBC1966) (Rantakallio *et al.*, 1969) with 16 traits having sample sizes between 4736 and 5025 individuals (Supplementary Table S1) and preprocessed by Tukiainen *et al.* (2014). We analyzed all 120 pairs of traits using both the complete (*t*_*i*_ = *t*_*d*_ = 0) and an approximate versions (*t*_*i*_ = *t*_*d*_ = 200) of biMM and compared with GCTA 1.25.3, GEMMA 0.94.1 and BOLT-REML 2.2 with their default parameters.

## 3 Results

Figure 1 shows that the complete and approximate versions of biMM are very similar across the 120 pairs of traits. Table 1 shows that the approximate version is much faster than either the complete version or any other software package tested. Detailed results are in Supplementary Figures S1-S4. In short, biMM and GEMMA gave essentially the same results and they were also similar to the results from GCTA and BOLT-REML.

**Table 1:** Cumulative run time over 120 trait pairs of Fig. 1. ‘Real’ is wall clock time. ‘CPU’ is total CPU time over all cores used. We used an Intel Quad-Core i7-3770 CPU @ 3.40GHz. biMM ran in R-3.3.1 with Intel Math Kernel Libarary.

|  | biMM approx | biMM compl | GEMMA | BOLT-REML | GCTA |
| --- | --- | --- | --- | --- | --- |
| Real (h) | 0.05 | 0.49 | 2.76 | 19.20 | 21.39 |
| CPU (h) | 0.07 | 1.49 | 2.76 | 19.20 | 21.39 |

**Figure 1:**
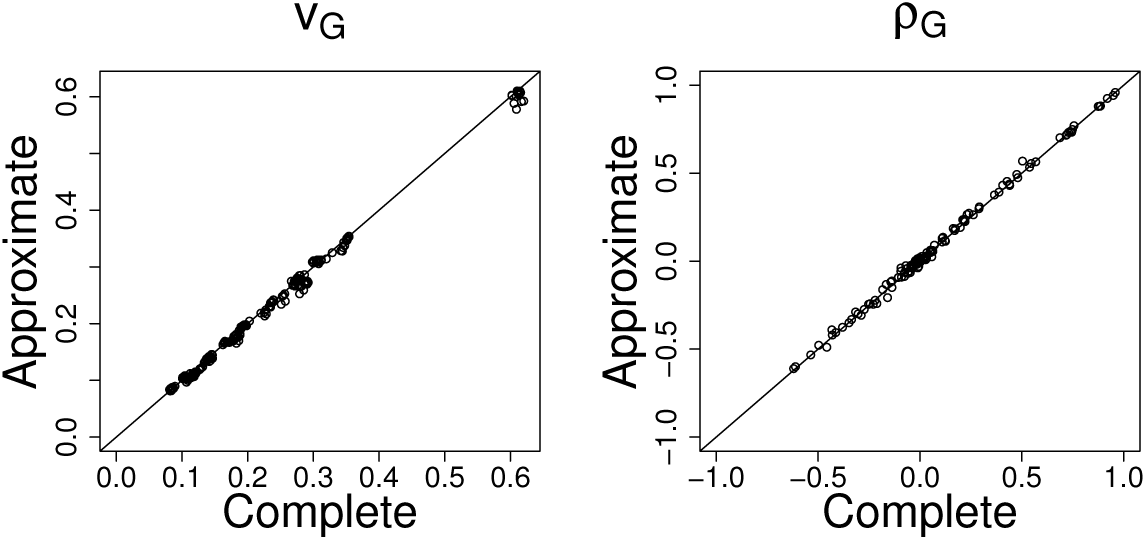
Comparing estimates for heritability (*V*_*G*_) and genetic correlation (*ρ*_*G*_) between an approximate (*t*_*i*_ = *t*_*d*_ = 200) and complete (*t*_*i*_ = *t*_*d*_ = 0) versions of biMM over 120 pairs of traits.

## Acknowledgements

This study made use of NFBC1966 data. We thank the late professor Paula Rantakallio (launch of NFBC1966), the participants in the 31yrs study and the NFBC project center.

